# artMAP: a user-friendly tool for mapping EMS-induced mutations in Arabidopsis

**DOI:** 10.1101/414433

**Authors:** Peter Javorka, Vivek Raxwal, Jan Najvarek, Karel Riha

**Author notes:** Equal contribution (Peter Javorka, Vivek Raxwal).

## Abstract

Mapping-by-sequencing is a rapid method for identifying both natural as well as induced variations in the genome. However, it requires extensive bioinformatics expertise along with the computational infrastructure to analyze the sequencing data and these requirements have limited its widespread adoption. In the current study, we develop an easy to use tool, artMAP, to discover ethyl methanesulfonate (EMS) induced mutations in the Arabidopsis genome. The artMAP pipeline consists of well-established tools including TrimGalore, BWA, BEDTools, SAMtools, and SnpEff which were integrated in a Docker container. artMAP provides a graphical user interface and can be run on a regular laptop and desktop, thereby limiting the bioinformatics expertise required. artMAP can process input sequencing files generated from single or paired-end sequencing. The results of the analysis are presented in interactive graphs which display the annotation details of each mutation. Due to its ease of use, artMAP made the identification of EMS-induced mutations in Arabidopsis possible with only a few mouse click. The source code of artMAP is available on Github (https://github.com/RihaLab/artMAP).

## Introduction

One of the key driving forces of evolution is *de novo* mutations that randomly occur in a genome, which may become fixed through natural selection or genetic drift. Natural genetic diversity can be used to identify genes responsible for phenotypes of interest either by standard and Quantitative Trait Locus (QTL) mapping or through random genome-wide integration of transgenes and transposons. Mutagenesis is then followed by the selection of mutant lines which exhibit the desired phenotype. Induced mutagenesis generates a much wider range of mutations than occur naturally, as many mutations would be selected against in natural populations. The key advantage of forward genetic screens over reverse genetic approaches, such as targeted gene knock-outs, is their ability to link biological functions to unknown genes in an unbiased manner. Furthermore, in contrast to knock-outs, irradiation and chemical mutagenesis produce a broad range of gene variants with different degrees of functionality, which can be instrumental for studying a gene’s regulation and its mechanism of action. For these reasons, forward genetic screens have been successfully applied in a number of model organisms to decipher the biological functions of many genes (Forsburg, 2001; Patton & Zon, 2001; Casselton & Zolan, 2002; Jorgensen & Mango, 2002; Page & Grossniklaus, 2002; St Johnston, 2002; Shuman & Silvahy, 2003; Grimm, 2004; Kile & Hilton, 2005; Candela & Hake, 2008).

While forward genetic screens are one of the most effective approaches for gene discovery, they still require a substantial time commitment and non-negligible monetary investments. While screening for a mutant line with a desired phenotype is often tedious, identification of the causative mutation is usually the main limiting factor in terms of resources, manpower, and time. This process usually involves the generation of mapping populations, which are used to associate a genomic region with the phenotype. The induced mutations in the associated region are then identified and causally linked to the phenotype by complementation tests or through the acquisition of independent alleles. In the pre-genomics era, mapping populations were derived from crosses with a genetically divergent strain that provided genetic markers for association mapping. Association mapping was used to identify the broader region of the genome and then followed by sequencing methods such as chromosome walking to identify the causative mutation. This approach is time-consuming and prone to many limitations, including the density of known polymorphisms in the divergent strain, introgression of unlinked phenotypic modulators, and distribution of meiotic crossovers. With the advent of Next Generation Sequencing (NGS) methodologies, many of these limitations were overcome by the direct sequencing of recombinant mapping populations. Because this approach identifies induced mutations genome-wide, mapping populations can be generated by back-crosses with parental strains using the *de novo* mutations as markers for association mapping (Hartwig *et al.*, 2012; Lindner *et al.*, 2012).

Forward genetic screens have been extensively used in Arabidopsis due to its well-annotated genome, self-pollination, and availability of genetic resources (Clouse *et al.*, 1996; Yin *et al.*, 2002; Manavella *et al.*, 2012; Berardini *et al.*, 2015). A genetic screen in Arabidopsis begins with the mutagenesis of seeds (M0), usually by EMS, followed by screening self-pollinated M_1_ or M_2_ plants for the phenotype of interest. Dominant mutations exhibit their phenotype in the M1 generation, whereas recessive mutations are scored in M_2_ (**Figure 1**). For both dominant and recessive mutations, M_2_ plants are either crossed with another Arabidopsis ecotype or back-crossed to the parental strain to produce recombinant mapping populations. The pool of plants displaying the desired phenotype is sequenced, providing the location of the associated genomic region and a set of candidate mutations. This approach greatly reduces the time and resources required to identify the causal mutation and also circumvents the dependence on genetic markers.

**Figure 1:**
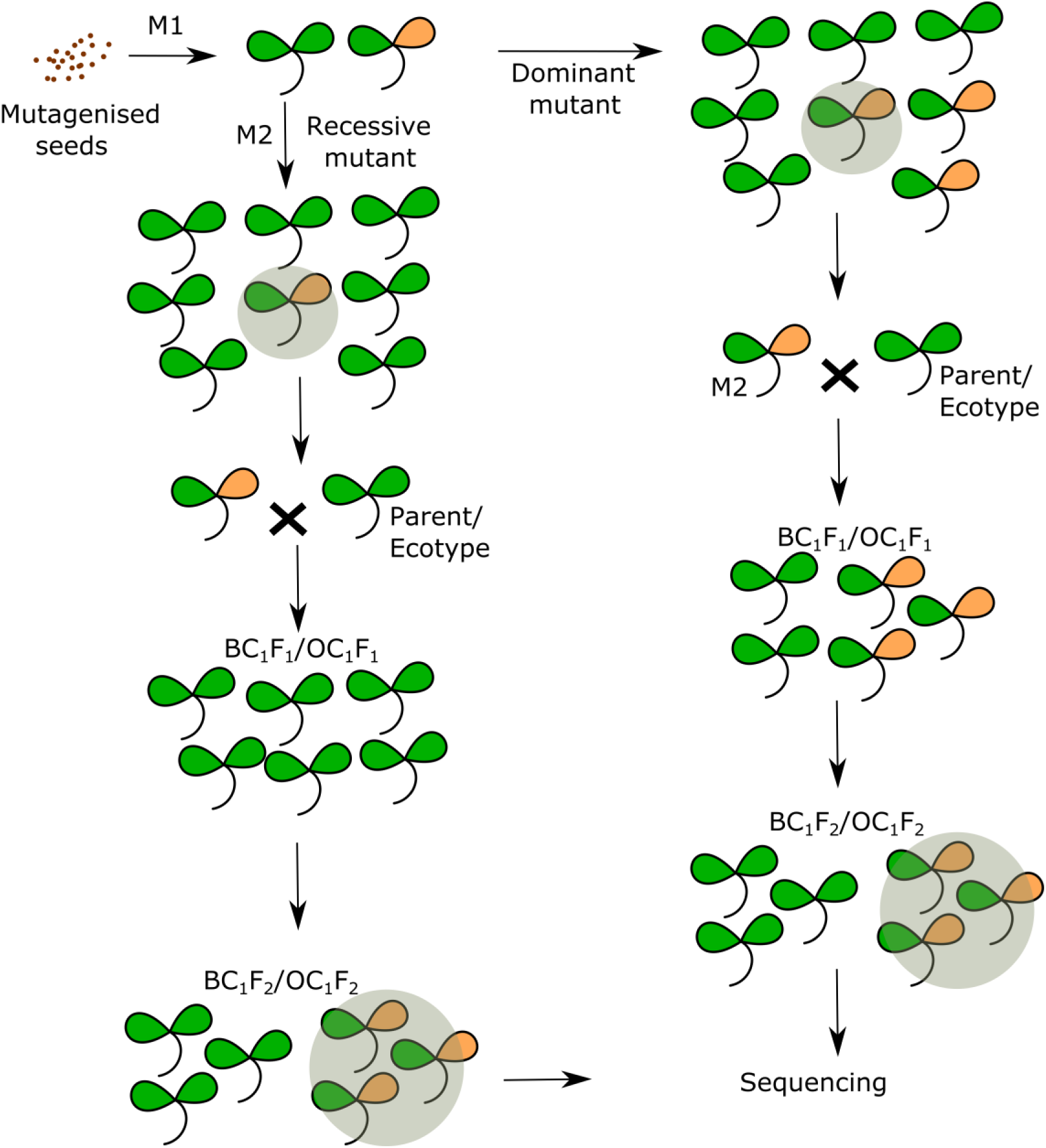
Schematic representation of forward genetic screens in Arabidopsis, showing the strategy for mapping dominant and recessive mutants

Two major bottlenecks in modern forward genetic screens are the high cost of NGS and the complexity of analyzing high throughput sequencing data. With the price of NGS falling continuously, data analysis remains the major bottleneck. While various pipelines have been developed to analyze sequencing data generated from forward genetic screens (Schneeberger *et al.*, 2009; Austin *et al.*, 2011; Minevich *et al.*, 2012; Wachsman *et al.*, 2017), they all require additional computational infrastructure along with bioinformatics expertise. Recently, SIMPLE was introduced to facilitate the analysis of forward genetic data, but even this method requires a certain level of bioinformatics understanding (Wachsman *et al.*, 2017). To date, there is no open source tool available which can be used by a biologist with no bioinformatics expertise. To fill this void, we developed an easy to use tool with a graphical user interface, artMAP, which can be used without any bioinformatics expertise to map EMS-induced mutations in Arabidopsis and asses their association with the desired phenotype.

## Result and Discussion

### Description of artMAP

The artMAP pipeline consists of various open sources tools integrated into a docker container (http://www.d>ocker.com/) to provide a graphical user interface (GUI) and the ability to run on all the three computer platforms (Windows/Mac/Linux). The pipeline is presented in **Figure 2**. Integrating any sequencing analysis pipeline into a single GUI faces several technical challenges as open source tools differ in their programming language, have multiple dependencies, may produce incompatibility issues when brought together. Moreover, the availability of a wide variety of sequencing platforms and format types increases the complexity. artMAP overcomes these issues by using five open source bioinformatics tools (SAMtools, BEDTools, BWA, Trimgalore, and SnpEff) along with in-house scripts to enable the analysis of all possible sequencing data types for EMS based genetic screens. Briefly, the artMAP pipeline consists of 6 steps: 1) pre-processing of the sequencing read files by Trimgalore (https://www.bioinformatics.babraham.ac.uk/projects/trim_galore/), >2) alignment of reads to the Arabidopsis genome by BWA (Li & Durbin, 2009), 3) post-processing of aligned reads by SAMtools (Li *et al.*, 2009), 4) identification of single nucleotide polymorphisms (SNPs) specific to mutant samples through the combined use of SAMtools and BEDTools suite (Quinlan & Hall, 2010), 5) Visualization of the SNPs, 6) annotation of SNPs by SnpEff (Cingolani *et al.*, 2012). Finally, artMAP provides a list of SNPs along with their allele frequency, depth, and annotation in a tab separated file.

**Figure 2:**
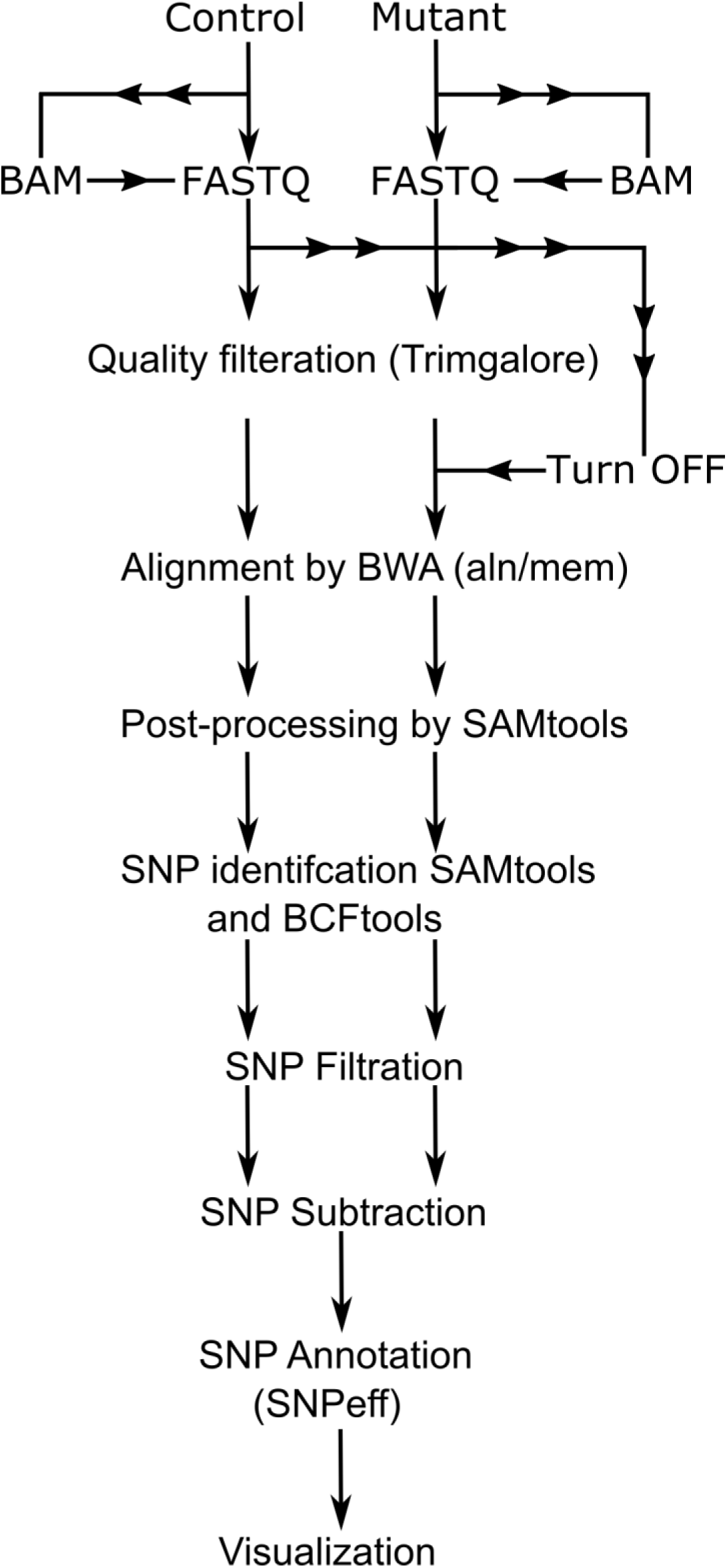
An outline of the artMAP pipeline showing

In forward genetic screens, whole genome sequencing read files are generated from two biological samples, one treated with EMS (mutant) and a control (usually the parent). The Illumina sequencing platform is preferred for re-sequencing applications, like mapping mutations, owing to its low error rate. An important parameter in NGS is the read length (in base pairs) and type of sequencing (single-end vs paired-end). Paired-end sequencing and a longer read length is often recommended for accurately mapping mutations, especially from the repetitive region of the genome. In most cases, however, single-end sequencing with high depth also enables mapping mutations with a single nucleotide resolution (James *et al.*, 2013; Wachsman *et al.*, 2017). artMAP can process both paired-and single-end sequencing reads regardless of their length. artMAP requires data in BAM or FASTQ formats, which are usually used by Illumina and other sequencing platforms. The data are first processed by Trimgalore, which remove sequencing adapters as well as bad quality sequences from the reads. Since Trimgalore process only FASTQ files, sequencing reads provided in the BAM format are converted to FASTQ using the bam2fastq function of BEDTools.

Next, high-quality sequencing reads from the mutant and parent samples are aligned to the Arabidopsis reference genome by BWA. The major advantage of BWA is the ability to align both short and long reads. artMAP include “BWA aln” as well as “BWA mem” for aligning shorter and longer reads, respectively, and artMAP can choose the appropriate aligner based on the length of the input reads. Also, as BWA requires an index of the genome to run, artMAP includes a pre-built BWA index of the Arabidopsis reference genome, eliminating the need to generate a genome index on every run. BWA not only reports the location of the sequencing reads but also records mismatches present between the genome and sequencing reads. This information is later exploited to identify SNPs. The results of the alignment from both control and mutant files are stored in a user-provided location in the SAM format. However, storing and downstream processing of these files are computationally inefficient as they require a large amount of operational memory (RAM) as well as storage space. To increase the computational efficiency, these aligned files are converted to a binary format (BAM). Handling and processing BAM files are computationally less demanding than SAM files. SAMtools is used to convert SAM to BAM files, which are then further sorted according to chromosome number and the genomic location of aligned reads (Li *et al.*, 2009). artMAP provides an additional option to control the removal of PCR duplicates from the control and mutant BAM files. This step is turned on by default but can be disabled if required. Control and mutant BAM files are indexed to prepare for SNP calling with SAMtools. The BAM alignment files generated in this step can be viewed in genome browsers such as IGV (Robinson *et al.*, 2011) for further analysis.

SNPs are identified from both the control and mutant BAM files generated in the previous step. These SNPs includes all single nucleotide changes present in the control and mutant samples. As EMS induces G to A or C to T transitions in DNA, artMAP only retains these SNPs. EMS induces a plethora of mutations in the genome, including both causative and non-causative SNPs. In the mapping population, the frequency of causative SNPs along with the surrounding linked SNPs is higher (approaching 100 % for a recessive mutation) compared to non-causative SNPs. artMAP applies two filter criteria to remove background polymorphisms that may occur as technical errors. First, artMAP removes SNPs with a frequency lower than 30%. Second, it applies a depth filter to retain SNPs with a sequencing depth between 10-100X. Since these filters can greatly affect the final outcome of the analysis, they can be changed in additional settings.

Next, to identify the region of the genome associated with the phenotype of interest, artMAP compares the SNPs present in the control and mutant sample and extracts SNPs exclusive to the mutant. These SNPs are then annotated using SnpEff (Cingolani *et al.*, 2012) to describe their impact on gene structure and amino acid changes. Nonsense mutations producing a stop codon are considered high impact SNPs. Further details regarding the SnpEff annotations can be obtained at http://snpeff.sourceforge.net/SnpEff.html#intro. artMAP displays the final results as a graph, where the frequency and position of each SNP are plotted along each Arabidopsis chromosome. These graphs can be zoomed in and can also be saved. Hovering the cursor over SNP reveals key information such as the location, frequency, affected gene, protein and DNA level changes, and the predicted the impact of the SNP. This visualization of the data facilitates a rapid assessment of the results and identification of the region associated with the phenotype.

Finally, artMAP provides results as a tab-delimited file with information containing the location of each SNP (chromosome number and position), reference base, mutated base, coverage over the base (depth), frequency, gene identity, and effect on protein change if any. Based on the graph and tab-delimited file, a user can identify the putative candidate gene for further testing. artMAP also produces the raw file at each stage of the pipeline, in case it is required. The detailed description of how to install and run artMAP is provided in supplemented User Manual.

### Implementation of artMAP to map EMS-induced mutations

First, we assessed the feasibility of mapping recessive mutations with artMAP. For this, we took data generated from the forward genetic screen for leaf hyponasty mutants (Allen *et al.*, 2013) where the recombinant mapping population (BC1F2) was produced by crossing the M2 plant with non-treated parent followed by one round of self-crossing. Since this screen is based on bulk segregation analysis of a recessive trait, the causal SNP should be present with a frequency of 100% and surrounded by a collection of linked, high-frequency SNPs. Unlinked SNPs should have a frequency of 50% as expected for the random inheritance of a heterozygous SNP within a population. We unambiguously identified a region linked to the phenotype on chromosome 3 with high-frequency mutations (**Figure 3**). This included a mutation in *HST1* that results in a stop codon at position 451 (Trp451*). This mutation was previously considered the causative mutation in this screen (Allen *et al.*, 2013).

**Figure 3:**
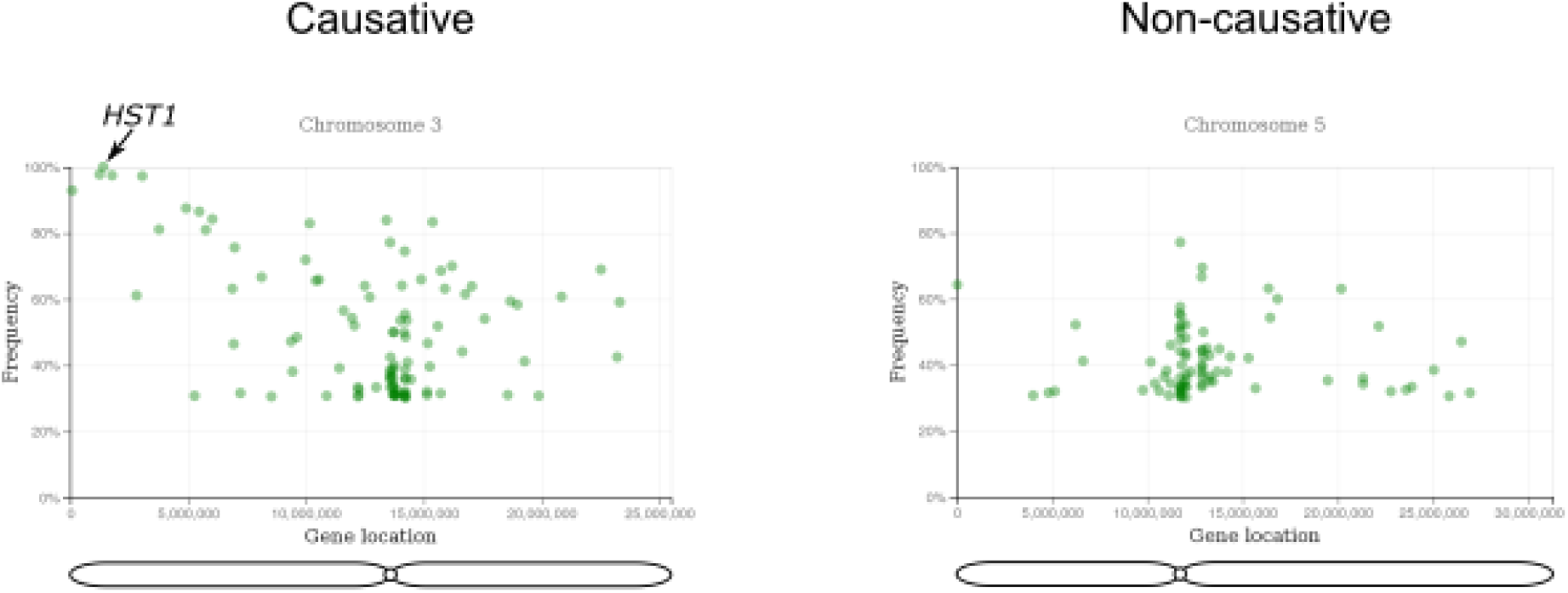
A representative figure of the example run showing the output of the artMAP analysis

Next, we assessed the performance of artMAP compared to a previously published pipeline. For this, we re-analyzed previously published datasets (Wachsman *et al.*, 2017) with SIMPLE (Wachsman *et al.*, 2017) as well as artMAP. This dataset included sequencing reads generated from single-as well as paired-end data. As expected, artMAP was able to accurately map the previously reported causative mutation or those identified by SIMPLE. The list of datasets used and results comparing artMAP and SIMPLE are presented in **Table 1**. It is important to note that while SIMPLE reports the list of likely candidate mutations, artMAP allows the user to interactively browse through graphs displaying the frequencies of individual mutations along the chromosomes, enabling the user to quickly define the linked region and assess whether a mutation may be causative. Also, artMAP displays annotation details and predicted SNP impact directly on the graph. This ability to easily manually assess mutations increases the probability of identifying the actual, causative mutation.

**Table 1:** List of datasets used to compare SIMPLE and artMAP

| Line | Reported gene | Reported mutation | Mapped by<br>SIMPLE | Mapped by<br>artMAP |
| --- | --- | --- | --- | --- |
| 300 | AT3G13870 | Ser584Phe | + | + |
| 300-4 | AT3G13870 | Ser584Phe | + | + |
| 300-7 | AT4G01800 | Arg752* | + | + |
| EMS608 | AT5G24630 | Gly324Glu | + | + |
| EMS633 | AT3G54660 | Ala264Thr | + | + |

## Conclusion

We have developed an interactive tool, artMAP, to map EMS-induced mutations in forward genetic screens in *Arabidopsis thaliana*. artMAP can easily be operated by researchers without any prior expertise in bioinformatics and we demonstrate that the accuracy of artMAP is similar to standard bioinformatics pipelines used to map EMS-induced mutations. It can be run on regular desktops or laptops and does not require extra computational infrastructure. Thus, artMAP greatly facilitates the identification of new mutations in forward genetic screens in Arabidopsis, and this tool can easily be adapted for other organisms, if needed.

## Acknowledgment

The authors are grateful for financial support by the Ministry of Education, Youth and Sports of the Czech Republic, European Regional Development Fund-Project "REMAP“ (No. CZ.02.1.01/0.0/0.0/15_003/0000479) and by the Czech Science Foundation (16-18578S).

